# Cell-type specific induction of cyclo-oxygenase-2 in layer II/III prefrontal cortical neurons mediates stress-induced anxiety phenotypes in mice

**DOI:** 10.1101/2021.10.10.463815

**Authors:** Robert J. Fenster, Kenneth McCullough, Sergey Naumenko, Andrew Thompson, Claudia Klengel, Allison Rodgers, Joy Otten, Dan Shu, Niki Harris, Torsten Klengel, Kerry J. Ressler

## Abstract

The ability of the medial prefrontal cortex (mPFC) to exert top-down control of behavior is affected by stress. The molecular response of mPFC to stress is incompletely understood, however, in part because of the region’s cellular heterogeneity. Here we used single nucleus RNA sequencing (snRNAseq) to map specific molecular cell types within the mPFC and to detect cell-type specific transcriptional changes to foot-shock stress. We identified *Ptgs2*, encoding cyclo-oxygenase 2, as an important candidate that is upregulated in layer II/III excitatory neurons after stress. Specifically, *Ptgs2* was transiently upregulated with shock-induced fear learning and fear expression, along with *Bdnf, Nptx2*, and *Lingo1*, in a layer II/III neuronal population marked by the neuronal excitatory gene *Slc17a7* and cell-type specific neuropeptide *Penk*. These dynamic cell-type specific expression patterns identified with snRNAseq were validated with quantitative fluorescent *in situ* hybridization. Using a pharmacological approach, we found that systemic lumiracoxib, a selective Ptgs2-inhibitor, led to a significant reduction in fear expression. Furthermore, genetic ablation of *Ptgs2* in excitatory *Camk2a*-expressing neurons led to reduced stress-induced anxiety-like behaviors. Together these findings suggest that *Ptgs2* is expressed in a dynamic, cell-type specific way in Layer II/III *Penk*+ neurons in mPFC, and that its role in prostaglandin and /or endocannabinoid regulation within these neurons may be an important mediator of stress-related behavior.

## Introduction

Understanding the role of cortical control over subcortical-driven emotional behavior is critical for progress in a wide range of psychiatric disorders from depression to PTSD[1]. Specifically, the medial prefrontal cortex (mPFC) has been implicated as being crucially important for both the development of and recovery from PTSD in humans and for the regulation and extinction of fear memories in rodent models[2-4]. Across most psychiatric disorders, stress can exert persistent negative effects and has been shown to lead to molecular, structural, and functional changes to the mPFC[5-8]. Because of the wide-ranging and pervasive impacts of stress on mental illness, understanding the molecular impact of stress on the mPFC is of great potential therapeutic interest.

Despite our knowledge of the significance of the mPFC in stress responsivity and the development of PTSD, we currently lack any specific pharmacological targets for stress-related disorders in this region. Our progress in the development of stress-related therapeutics that target the mPFC relates in part to the enormous complexity of the structure. The rodent mPFC is a highly organized laminar structure composed of many cell classes—excitatory projection neurons, inhibitory interneurons, glia, microglia, and endothelial cells—each with many different molecular subtypes[9-11]. For many years, studies of molecular changes in brain, and cortex more specifically, relied upon bulk RNA sequencing of transcripts from tissue homogenates. These studies allowed for the identification of important molecular regulators of the stress response[12-15]. However, such approaches are likely to only scratch the surface of the complexity of the molecular responses. They may obscure cell-type specific changes due to a lack of sensitivity and due to variation in function across cells that are anatomically intermixed. Notably, recent advances in cell-type specific and single cell RNA sequencing technologies[16,17] offer opportunities to detect molecular changes in specific cell types that might not have been discovered with earlier methodologies.

In this study, we use single nucleus RNA sequencing (snRNAseq) to identify cell-type specific molecular responses to an auditory Pavlovian fear conditioning paradigm after fear training and expression. We identify *Ptgs2*, the gene encoding Cyclo-oxygenase-2, as being increased in layer II/III mPFC neurons after fear conditioning and fear expression. Further we demonstrate that Ptgs2 inhibition leads to anxiolytic phenotypes.

## Results

### snRNAseq of mPFC from home cage, fear conditioned and fear expressing mice

To understand molecular mechanisms of fear regulation, our initial goal was to identify differential cell-type specific gene expression in mouse medial prefrontal cortex (mPFC), a brain region known to regulate fear learning, expression, and inhibition. We sought to identify differential transcriptional signatures during the consolidation phase of fear learning, or after fear expression, compared to home cage controls. Thus, we collected RNA from fresh-frozen mPFC punches (1mm) from naïve, adult C57Bl/6J mice and from mice that were sacrificed two hours after Pavlovian auditory fear conditioning or two hours after fear expression. We then performed single nuclear RNA sequencing (snRNAseq) and compared the transcriptional profiles across mice from these three conditions.

We profiled a total of 84,330 nuclei that passed quality control (**Fig. 1A**). Sequenced nuclei that passed filtering had a mean of 1,665 unique molecular identifiers (UMIs)/nucleus and a mean of 1,220 detectable expressed genes/nucleus (**Fig. 2)**. To generate specific cell-type specific identifiers, nuclei across all experimental conditions were initially clustered together. Using the Uniform Manifold Approximation and Projection (UMAP) method for dimensionality reduction, we identified a total of 26 clusters (**Fig.1B**). Identified clusters were distributed evenly across conditions, suggesting limited batch effects and indicating that cell-type identifiers were not altered by behavioral condition.

**Figure 1:**
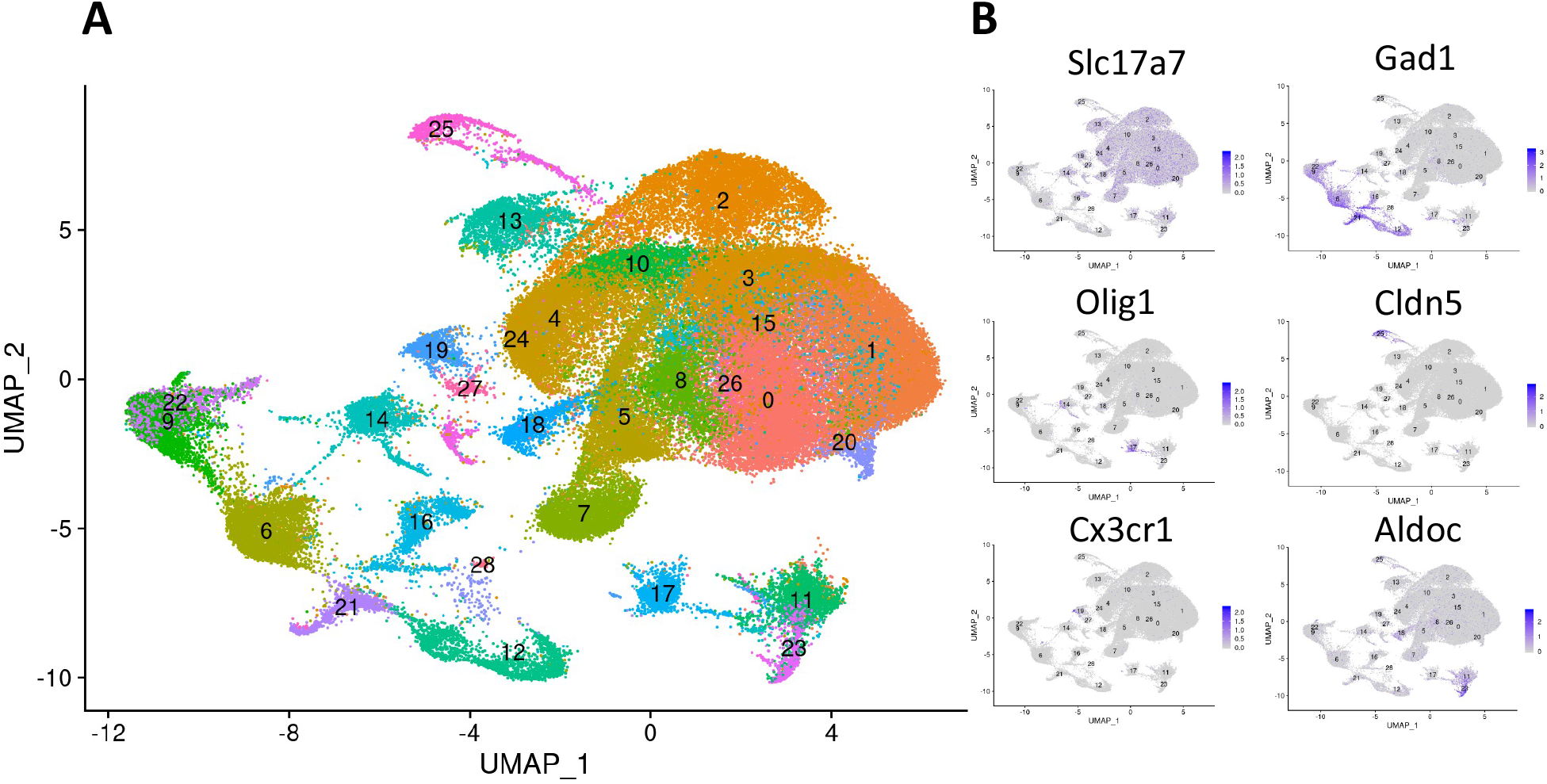
UMAP plot of mPFC nuclei. A) nuclei across all conditions were clustered together and visualized in this UMAP plot. B) Markers for excitatory neurons (Slc17a7), inhibitory neurons (Gad1), oligodendrocytes (Olig1), endothelial cells (Cldn5), microglia (Cx3cr1), and astrocytes (Aldoc) are plotted on the UMAP, marking distinct territories.

**Figure 2:**
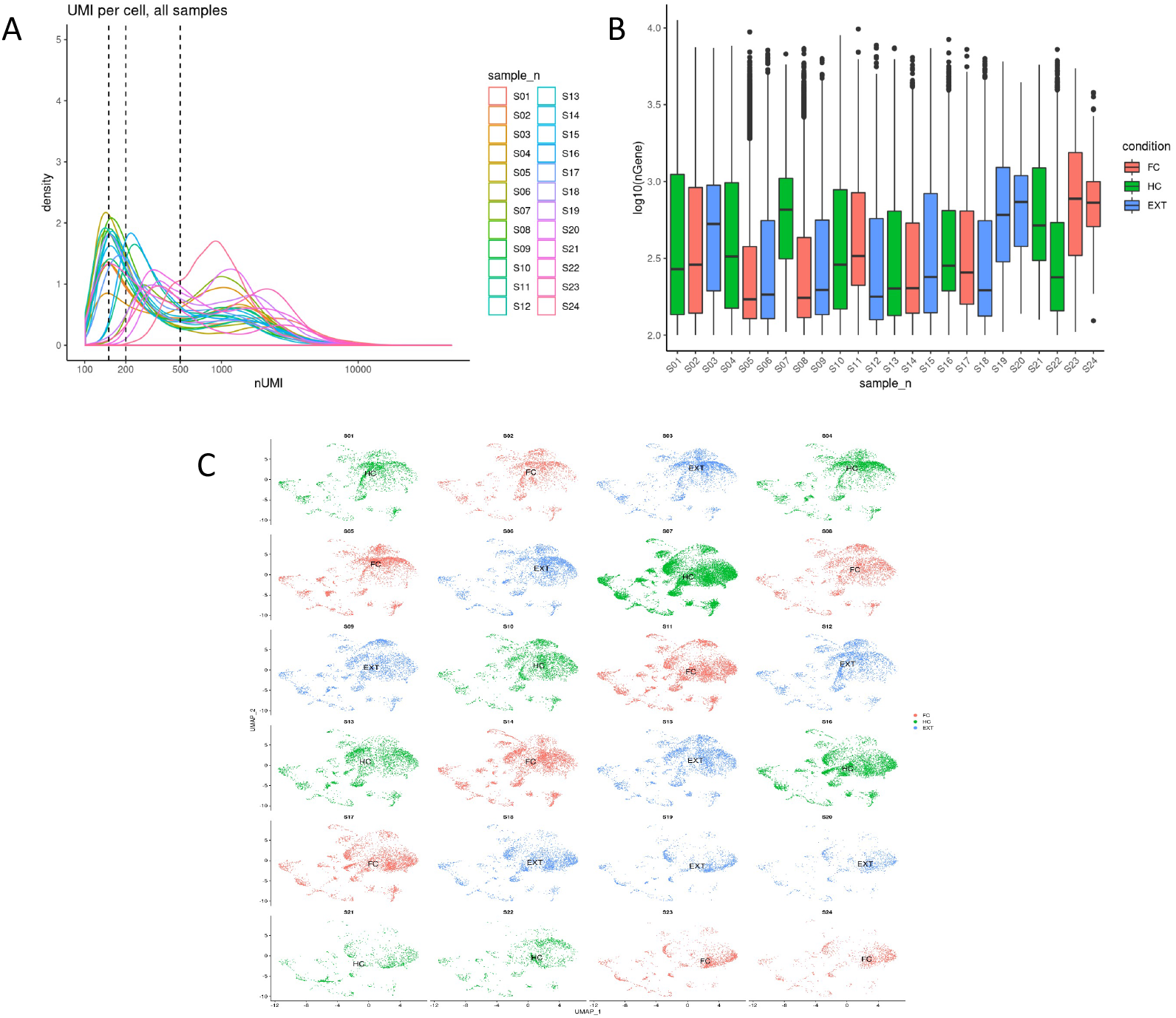
Quality control metrics for snRNAseq data. A) Histogram describing the spread of UMIs/nucleus. B) Boxplot showing the range of the number of genes detected/nucleus across all samples. C) UMAP plots across all samples showing relative homogeneity of all clusters across samples.

Our data demonstrated discrete subgroupings of clusters in mouse mPFC that express known markers of excitatory neurons (*Slc17a7*), inhibitory neurons (*Gad1*), oligodendrocytes (*Olig1*), endothelial cells (*Cldn5*), microglia (*Cx3cr1*), and astrocytes (*Aldoc*) (**Fig. 1B**). Nuclei were sub-clustered by expression of *Gad1* and *Slc17a7*, to detect additional cell-types within the data. These sub-clusters were used to compute differential expression analyses. The inhibitory subcluster taxonomy includes markers for cell types known to arise out of the caudal ganglionic eminence (CGE), *Lamp5* and *Vip*, and medial ganglionic eminence (MGE), including *Sst* and *Pvalb* [10,18]. As previously observed in other single-cell datasets from mouse cortex[10], there is more continuity between glutamatergic cell-types, with cortical depth driving much of the variability between clusters. Clusters contained known layer-specific markers, including *Wfs1* (layer II/III), *Penk* (layer II/III), *Cck* (layer IV), *Fezf2, Pou3f1*, and *Bcl11b* (layer V), and *Rprm* (layer VI).

### Differential expression analysis

We next performed differential expression analysis across conditions using the Seurat package. A full list of differentially expressed genes is available upon request. We prioritized follow-up of cell-types that exhibit changes in genes known to be involved in neural plasticity mechanisms. One cluster of excitatory neurons most strongly marked by neural plasticity activation was the *Penk*-expressing cluster. In the Allen Brain Atlas *in situ* hybridization database[19], *Penk* is expressed most strongly in a subset of layer II/III neurons in the mPFC. Our data showed that with fear behaviors, the *Penk+* cells were marked by dynamically upregulated *Bdnf*, a neural plasticity gene that has reproducibly been shown to be necessary in fear learning and expression processes [20-24]. In addition to *Bdnf*, this cluster exhibited changes to *Nptx2*, which is a known glutamatergic synapse plasticity-regulator[25-27], as well as *Lingo1* and non-coding RNA *Malat1*, which have also been implicated in synaptic plasticity[28,29] (**Fig. 3**). Interestingly, the gene that was most significantly changed in this cell-type was *Ptgs2*. Ptgs2 was increased significantly in the fear expression condition and trended toward increase in the fear conditioning group (**Fig. 3**). Using FISH, we confirmed that *Ptgs2* and *Bdnf* are largely co-expressed in layer II/III cells within mouse mPFC (**Fig. 4**) and are present in both prelimbic (PL) and infralimbic (IL) cortex.

**Figure 3:**
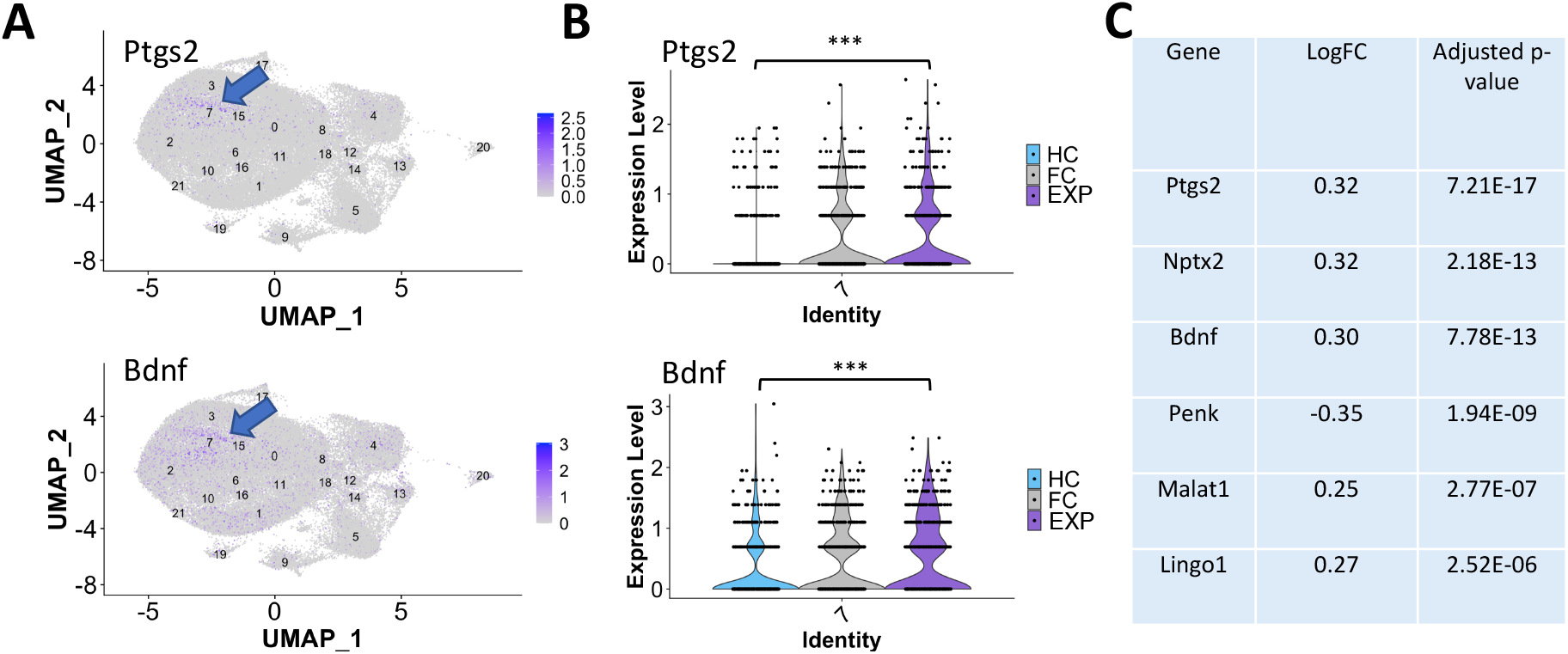
Ptgs2 and Bdnf are co-expressed in subset of layer II/III neurons. A) UMAP plot of excitatory neurons showing co-expression of Bdnf and Ptgs2 in cluster 7 along with violin plot (B) demonstrating increase in Ptgs2 and Bdnf with fear expression. C) table with differentially expressed genes in cluster 7.

**Figure 4:**
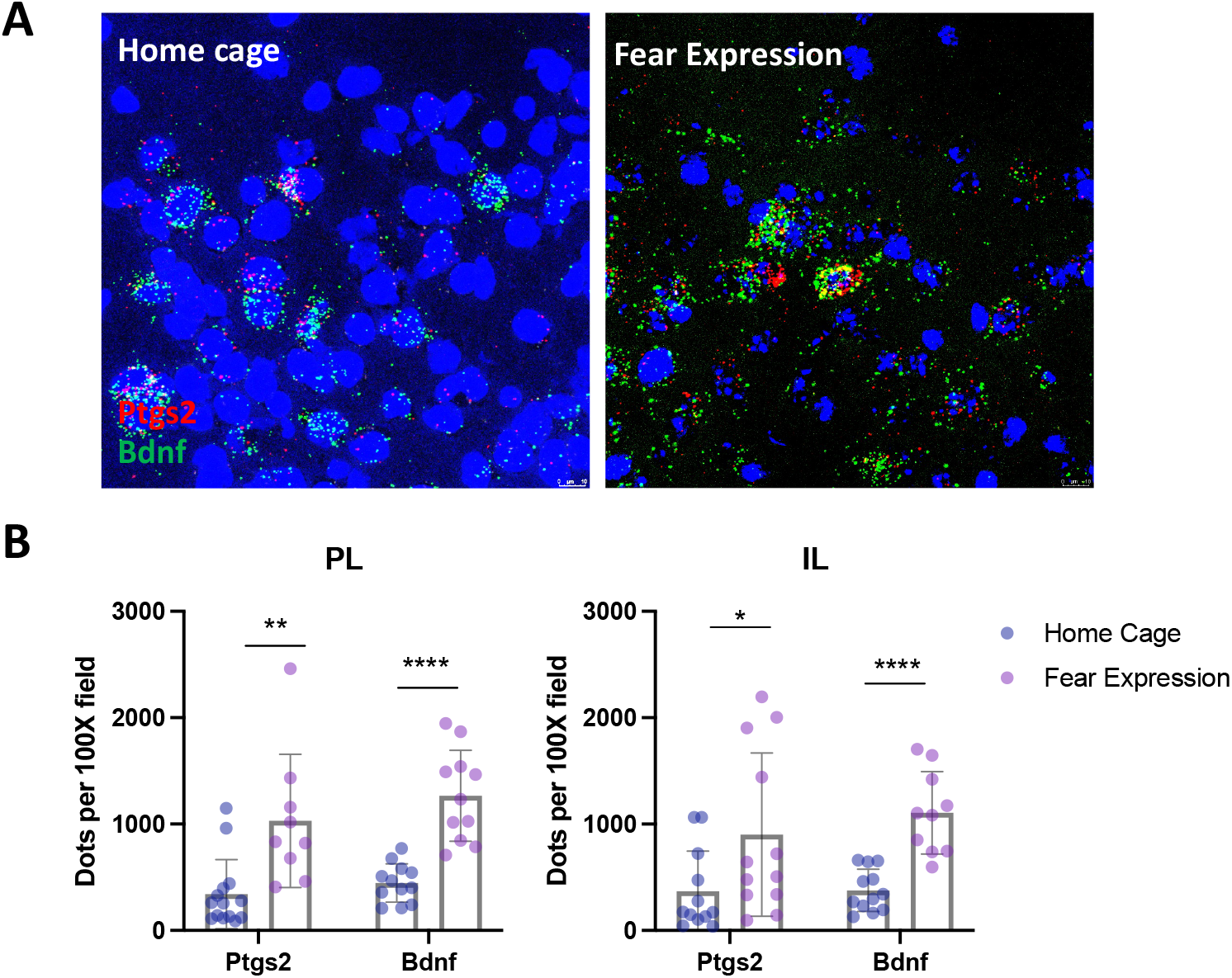
Ptgs2 is increased in mPFC after fear expression. A) left, Representative 100X confocal images from FISH to Ptgs2 and Bdnf in mPFC of home cage and fear expression groups. right, B) Quantification of increase in Ptgs2 and Bdnf in infralimbic (IL) and prelimbic (PL) cortices, n=2 sections from n=3 animals per group). Columns show mean ± S.E.M. Significance from unpaired t-test, p-values, left to right p=0.002, p=0.04, p<0.0001, and p<0.0001.

### Inhibition of Ptgs2 leads to anxiolysis

*Ptgs2* (also known as Cyclo-oxygenase-2 or Cox-2), encodes an inducible enzyme that has two known functions within the brain: it catalyzes the key step of prostaglandin synthesis and also metabolizes endogenous endocannabinoids, anandamide (AEA) and 2-arachidonoylglycerol (2-AG)[30]. *Ptgs2*/*Cox-2* has been implicated in the brain’s adaptations to chronic stress[31], as well as in the storage of long-term memory[32]. Recently, decreases in the levels of 2-AG within layer II/III PL neurons has also been shown to enhance reciprocal connectivity between BLA and PL, leading to anxiety-like behaviors after stress [33].

Given these prior findings, we hypothesized that *Ptgs2* upregulation in *Penk*-expressing layer II/III neurons might mediate the stress-induced anxiety-like behaviors after fear conditioning. To address this directly, we injected mice systemically with lumiracoxib, a selective Ptgs2-inhibitor, or vehicle, 30 minutes prior to fear expression. Lumiracoxib caused a significant reduction in fear expression compared to vehicle (**Fig. 5A**, Two-way ANOVA, Drug treatment, F = 5.877, p-value = 0.0275).

**Figure 5:**
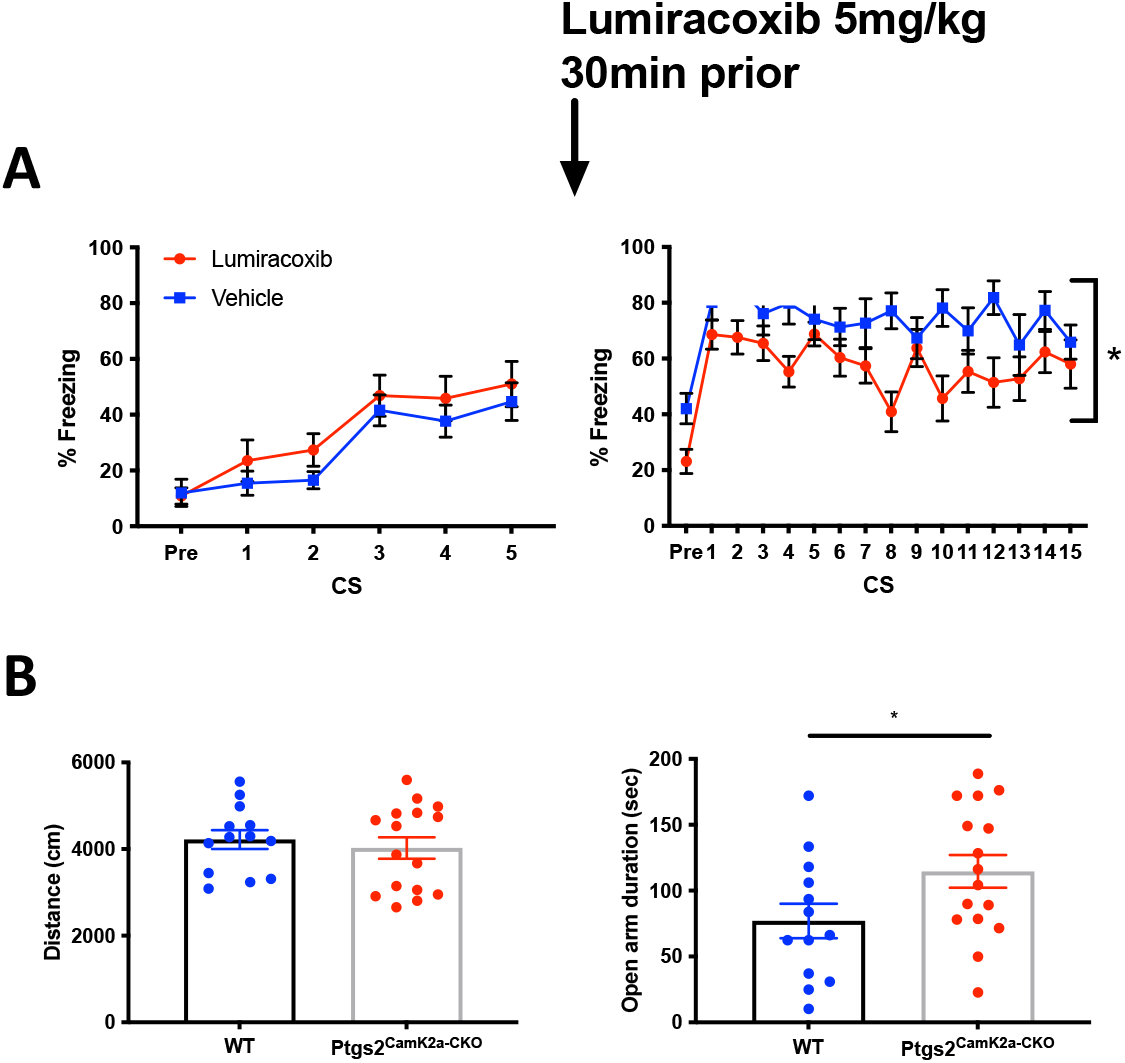
Inhibition of Ptgs2 blunts stress-induced anxiety-like behaviors. A) adult male C57Bl6/J mice were subjected to fear conditioning, and then, 24 hours later, underwent fear expression/fear extinction training in a novel context. Mice received systemic lumiracoxib injections 30 minutes prior to fear expression. Two-way ANOVA, Drug treatment, F = 5.877, p-value = 0.0275. B) male and female adult mice, homozygous for the floxed-Ptgs2 allele, either positive for CamK2a-Cre (Ptgs2^Camk2a-CKO^, n=16) or negative for the allele, WT (n=13), were tested in the elevated plus maze and open field. Ptgs2^Camk2a-CKO^ demonstrated an anxiolytic phenotype in the EPM (Unpaired two-tailed t-test, p=0.0471), but no locomotor differences in the open field.

To confirm that this pharmacological effect was not due to off-target effects, we performed a genetic ablation of Ptgs2 in all excitatory neurons by crossing a floxed-Ptgs2 mouse line[34] to Camk2a-Cre. We saw no effects on fear conditioning or fear expression with this manipulation (data not shown); however, we observed that mice that lost Ptgs2 in excitatory neurons exhibited an anxiolytic phenotype (less time in open arms) in the elevated plus maze (**Fig. 5B**, Students Two-Tailed t-test, p = 0.0471).

## Discussion

We generated an snRNAseq dataset of mouse mPFC to dissect cellular specificity and heterogeneity at baseline and to determine differential dynamic transcriptional profiles after fear conditioning and fear expression. Our data suggest an increase to the activity-dependent gene *Ptgs2/Cox-2* along with *Bdnf* in layer II/III *Penk*-expressing excitatory neurons in the mPFC following fear expression. Inhibition of Cox-2 with the selective inhibitor lumiracoxib leads to a decrease in fear expression behaviors, and genetic ablation of *Ptgs2* in Camk2a-expressing cells leads to an anxiolytic phenotype in the elevated plus maze. Our results are consistent with previous studies that have shown lumiracoxib to reduce anxiety-related behaviors after stress[35] and that constitutive knockout of Ptgs2 results in a mild anxiolytic phenotype[36]. Here, we specifically implicate stress-responsive changes within specific cell populations in mPFC as a potential mechanism for this effect.

Recent work has implicated the loss of 2-AG signaling in layer II/III mPFC neurons as a mechanism for strengthening connectivity between PFC and BLA, resulting in increased anxiety-like behaviors[33]. Ptgs2 has been shown to degrade endocannabinoids *in vivo*, and could play a role in mediating the loss of 2-AG: although MAGL is thought to be the primary enzyme responsible for 2-AG metabolism, there is a suggestion that Ptgs2 may metabolize it *in vivo* [36]. It is also possible that Cox-2 prostaglandin synthesis capacity is responsible for mediating its role in stress reactivity.

There have been reports that *Ptgs2* overexpression can drive *Bdnf* expression through PKA signaling[37], but it is unclear in this cell type whether Bdnf and Ptgs2 are co-regulated by upstream factors or whether one induces the other. Future work to elucidate the signal transduction cascades, as well as the transcriptional regulation of *Ptgs2* is warranted.

In summary, we find a cell-type specific upregulation of Ptgs2/Cox-2 in mouse PFC after foot shock stress and show that Cox-2 inhibition blunts stress-induced anxiety-like behaviors in mice. Although Cox-2 inhibitors were pulled from the market due to concerns for cardiovascular risk over chronic use, they could be a potential novel therapeutic for stress-induced anxiety, as has been previously proposed[30]. Further understanding of cell-type specific regulation of learning and stress-related molecular pathways offers significant promise towards advancing knowledge of, and therapeutic approaches to, brain mechanisms of behavior and behavioral disorders.

## Materials and Methods

### Animal Husbandry/Mouse Strains

All animals were kept on a 12 hour light-dark cycle (7:00 am -07:00 pm) and given *ad libitum* access to food and water. Mouse experiments were performed in accordance with the McLean Hospital Institutional Animal Care and Use of Laboratory Animals Committee. C57Bl/6J male mice were purchased from the Jackson Laboratory (Bar Harbor, ME). Conditional *Ptgs2* knockout mice were generated by breeding floxed-Ptgs2 mice[34] (Jackson Laboratories, strain 030785) and CamK2a-Cre mice (Jackson Laboratories). All mice were behaviorally tested between 2-5 months of age. Lumiracoxib was purchased from Abcam (Cambridge, MA).

### Behavior

#### Fear conditioning

Mice were habituated to the fear conditioning chamber (Med Associates, St. Albans, VT) for 20 minutes each day prior to testing. On the day of testing, mice were exposed to five tone-shock pairings (pre-CS period: 180 s, CS: 30s 6000 Hz, 65–70 db, co-terminating shock (US): 0.5 s, 0.7 mA, variable ITI). The fear expression group was returned to the animal care facility, while the fear conditioned group was left in the holding room until sacrifice, 2 hours after training. The next day, the fear expression group was placed in a novel context and presented with 30 CS presentations without any US reinforcement (pre-CS period: 180 s, CS: 30 s 6000 Hz, 65–70 db, ITI between CS’s: 60 s). Following fear expression, mice were kept in the holding room until sacrifice, 2 hours later. Freezing was measured using FreezeFrame5 software (Actimetrics, Wilmette, IL).

#### Elevated plus maze

For the elevated plus maze test, animals were placed on the open arm, facing the center. Apparatus had arms measuring 50 cm tip to tip and was placed in a dimly lit room. Mice were allowed to freely explore for 10 min, while behavior was recorded. Time spent in open arms, closed arms, and center as calculated using Ethovision software.

#### Tissue preparation

After sacrifice, brains were rapidly dissected and frozen on dry ice powder. Brains were kept stored at −80 C until punching, when they were sectioned and punched on a microtome. Punches were stored at −80 C until the day of nuclear isolation and encapsulation.

#### Single nucleus RNA sequencing

Nuclei were isolated as in previous studies[38], with minor modifications. Punches were placed into buffer HB (0.25 M sucrose, 25 mM KCl, 5 mM MgCl_2_, 20 mM Tricine-KOH pH 7.8, 1 mM DTT, 0.15 mM spermine, 0.5mM spermidine, protease inhibitors) and placed in a Dounce homogenizer for 10 strokes each with loose and tight pestles. A 5% IGEPAL solution was added to a final concentration of 0.16% followed by five additional dounce strokes, then the lysate was filtered through a 40-μm strainer. Nuclei were mixed with and equal volume o 50% iodixanol and then layered on top of an iodixanol gradient of 40% and 30% layers in a 2 mL dolphin microcentrifigue tube. Nuclei were spun by centrifuging at 10,000 x g for 4 min at 4°C and then collected by aspiration at the interface of the 30% and 40% iodixanol layers. Nuclear concentration and prep quality were ascertained by loading on a hemocytometer and were diluted to a concentration of 80-100K and 15% iodixanol with Buffer HB prior to loading on InDrops platform. Single-nuclei suspensions were encapsulated into droplets, lysed, and the RNA within each droplet was reverse-transcribed using unique nucleotide barcode as described previously[17]. Approximately 3000 cells were processed per library and sequenced on Illumina NovaSeq S2 chips (at a density of approximately 20,000 reads/nucleus).

#### Bioinformatics

We used bcbio-nextgen to process data starting from Illumina fastq files to counts. Namely, we parsed barcode information, demultiplexed reads by samples, filtered erroneous barcodes, and calculated sample-cell-barcode counts with umis. Before counting, we created a pseudoalignment with rapmap. All steps are described in detail in the documentation (https://bcbio-nextgen.readthedocs.io/en/latest/contents/single_cell.html#st…). We created a Seurat object using the count matrix and continued analysis (quality control, clustering, markers, differential expression) in Seurat using R (https://www.r-project.org/) and r-studio (https://rstudio.com/). To create quality control plots we used ggplot2, tidyverse (https://www.tidyverse.org/) and knitr (https://yihui.org/knitr/). We used mm10 mouse genome reference (https://www.ncbi.nlm.nih.gov/assembly/GCF_000001635.20/) and ensembl-94 annotation of genes (ftp://ftp.ensembl.org/pub/release-94/gtf/mus_musculus).

#### FISH

We used RNAscope Multiplex Fluorescent Assay (ACDBio, Newark, CA) to assay for Ptgs2 (316621-C2) and Bdnf (424821-C1). Quantification was performed using StarSearch algorithm from the Raj lab.

## Acknowledgments

This work was supported by NIH awards P50-MH115874, R01-MH108665, and the Frazier Institute at McLean Hospital. (KJR).

## References

1 Maddox SA, Hartmann J, Ross RA, Ressler KJ. Deconstructing the Gestalt: Mechanisms of Fear, Threat, and Trauma Memory Encoding. Neuron. 2019;102(1):60–74.

2 Fenster RJ, Lebois LAM, Ressler KJ, Suh J. Brain circuit dysfunction in post-traumatic stress disorder: from mouse to man. Nat Rev Neurosci. 2018;19(9):535–51.

3 Shin LM, Rauch SL, Pitman RK. Amygdala, medial prefrontal cortex, and hippocampal function in PTSD. Annals of the New York Academy of Sciences. 2006;1071:67–79.

4 Giustino TF, Maren S. The Role of the Medial Prefrontal Cortex in the Conditioning and Extinction of Fear. Front Behav Neurosci. 2015;9:298.

5 de Kloet ER, Joëls M, Holsboer F. Stress and the brain: from adaptation to disease. Nat Rev Neurosci. 2005;6(6):463–75.

6 Lupien SJ, McEwen BS, Gunnar MR, Heim C. Effects of stress throughout the lifespan on the brain, behaviour and cognition. Nat Rev Neurosci. 2009;10(6):434–45.

7 McEwen BS, Bowles NP, Gray JD, Hill MN, Hunter RG, Karatsoreos IN, et al. Mechanisms of stress in the brain. Nat Neurosci. 2015;18(10):1353–63.

8 Alexandra Kredlow M, Fenster RJ, Laurent ES, Ressler KJ, Phelps EA. Prefrontal cortex, amygdala, and threat processing: implications for PTSD. Neuropsychopharmacology. 2021.

9 Saunders A, Macosko EZ, Wysoker A, Goldman M, Krienen FM, de Rivera H, et al. Molecular Diversity and Specializations among the Cells of the Adult Mouse Brain. Cell. 2018;174(4):1015-30.e16.

10 Yao Z, Liu H, Xie F, Fischer S, Adkins RS, Aldridge AI, et al. A transcriptomic and epigenomic cell atlas of the mouse primary motor cortex. Nature. 2021;598(7879):103–10.

11 A multimodal cell census and atlas of the mammalian primary motor cortex. Nature. 2021;598(7879):86–102.

12 Choi DC, Maguschak KA, Ye K, Jang SW, Myers KM, Ressler KJ. Prelimbic cortical BDNF is required for memory of learned fear but not extinction or innate fear. Proc Natl Acad Sci U S A. 2010;107(6):2675–80.

13 Andero R, Ressler KJ. Fear extinction and BDNF: translating animal models of PTSD to the clinic. Genes Brain Behav. 2012;11(5):503–12.

14 Shansky RM, Hamo C, Hof PR, McEwen BS, Morrison JH. Stress-induced dendritic remodeling in the prefrontal cortex is circuit specific. Cereb Cortex. 2009;19(10):2479–84.

15 McEwen BS, Nasca C, Gray JD. Stress Effects on Neuronal Structure: Hippocampus, Amygdala, and Prefrontal Cortex. Neuropsychopharmacology. 2016;41(1):3–23.

16 Macosko EZ, Basu A, Satija R, Nemesh J, Shekhar K, Goldman M, et al. Highly Parallel Genome-wide Expression Profiling of Individual Cells Using Nanoliter Droplets. Cell. 2015;161(5):1202–14.

17 Klein AM, Mazutis L, Akartuna I, Tallapragada N, Veres A, Li V, et al. Droplet barcoding for single-cell transcriptomics applied to embryonic stem cells. Cell. 2015;161(5):1187–201.

18 Kepecs A, Fishell G. Interneuron cell types are fit to function. Nature. 2014;505(7483):318–26.

19 Lein ES, Hawrylycz MJ, Ao N, Ayres M, Bensinger A, Bernard A, et al. Genome-wide atlas of gene expression in the adult mouse brain. Nature. 2007;445(7124):168–76.

20 Sun W, Li X, An L. Distinct roles of prelimbic and infralimbic proBDNF in extinction of conditioned fear. Neuropharmacology. 2018;131:11–19.

21 Rosas-Vidal LE, Do-Monte FH, Sotres-Bayon F, Quirk GJ. Hippocampal--prefrontal BDNF and memory for fear extinction. Neuropsychopharmacology. 2014;39(9):2161–9.

22 Radiske A, Rossato JI, Köhler CA, Gonzalez MC, Medina JH, Cammarota M. Requirement for BDNF in the reconsolidation of fear extinction. J Neurosci. 2015;35(16):6570–4.

23 Peters J, Dieppa-Perea LM, Melendez LM, Quirk GJ. Induction of fear extinction with hippocampal-infralimbic BDNF. Science. 2010;328(5983):1288–90.

24 Bredy TW, Wu H, Crego C, Zellhoefer J, Sun YE, Barad M. Histone modifications around individual BDNF gene promoters in prefrontal cortex are associated with extinction of conditioned fear. Learn Mem. 2007;14(4):268–76.

25 Wang Z, Jin T, Le Q, Liu C, Wang X, Wang F, et al. Retrieval-Driven Hippocampal NPTX2 Plasticity Facilitates the Extinction of Cocaine-Associated Context Memory. Biological psychiatry. 2020;87(11):979–91.

26 Mariga A, Glaser J, Mathias L, Xu D, Xiao M, Worley P, et al. Definition of a Bidirectional Activity-Dependent Pathway Involving BDNF and Narp. Cell reports. 2015;13(9):1747–56.

27 Elbaz I, Lerer-Goldshtein T, Okamoto H, Appelbaum L. Reduced synaptic density and deficient locomotor response in neuronal activity-regulated pentraxin 2a mutant zebrafish. FASEB journal : official publication of the Federation of American Societies for Experimental Biology. 2015;29(4):1220–34.

28 Trifunovski A, Josephson A, Ringman A, Brené S, Spenger C, Olson L. Neuronal activity-induced regulation of Lingo-1. Neuroreport. 2004;15(15):2397–400.

29 Lipovich L, Dachet F, Cai J, Bagla S, Balan K, Jia H, et al. Activity-dependent human brain coding/noncoding gene regulatory networks. Genetics. 2012;192(3):1133–48.

30 Patel S, Hill MN, Cheer JF, Wotjak CT, Holmes A. The endocannabinoid system as a target for novel anxiolytic drugs. Neurosci Biobehav Rev. 2017;76(Pt A):56-66.

31 Morgan AJ, Kingsley PJ, Mitchener MM, Altemus M, Patrick TA, Gaulden AD, et al. Detection of Cyclooxygenase-2-Derived Oxygenation Products of the Endogenous Cannabinoid 2-Arachidonoylglycerol in Mouse Brain. ACS chemical neuroscience. 2018;9(7):1552–59.

32 Teather LA, Packard MG, Bazan NG. Post-training cyclooxygenase-2 (COX-2) inhibition impairs memory consolidation. Learning & memory (Cold Spring Harbor, NY). 2002;9(1):41–7.

33 Marcus DJ, Bedse G, Gaulden AD, Ryan JD, Kondev V, Winters ND, et al. Endocannabinoid Signaling Collapse Mediates Stress-Induced Amygdalo-Cortical Strengthening. Neuron. 2020;105(6):1062-76.e6.

34 Ishikawa TO, Herschman HR. Conditional knockout mouse for tissue-specific disruption of the cyclooxygenase-2 (Cox-2) gene. Genesis (New York, NY : 2000). 2006;44(3):143–9.

35 Morgan A, Gaulden A, Altemus M, Williford K, Centanni S, Winder D, et al. Cyclooxygenase-2 inhibition prevents stress induced amygdala activation and anxiety-like behavior. Brain Behav Immun. 2020;89:513–17.

36 Hermanson DJ, Hartley ND, Gamble-George J, Brown N, Shonesy BC, Kingsley PJ, et al. Substrate-selective COX-2 inhibition decreases anxiety via endocannabinoid activation. Nat Neurosci. 2013;16(9):1291–8.

37 Luo Y, Kuang S, Li H, Ran D, Yang J. cAMP/PKA-CREB-BDNF signaling pathway in hippocampus mediates cyclooxygenase 2-induced learning/memory deficits of rats subjected to chronic unpredictable mild stress. Oncotarget. 2017;8(22):35558–72.

38 Renthal W, Boxer LD, Hrvatin S, Li E, Silberfeld A, Nagy MA, et al. Characterization of human mosaic Rett syndrome brain tissue by single-nucleus RNA sequencing. Nat Neurosci. 2018;21(12):1670–79.

